# PccGEO: prior constraints conditioned genetic elements optimization

**DOI:** 10.1101/2021.11.08.467823

**Authors:** Hanwen Xu, Pengcheng Zhang, Haochen Wang, Lei Wei, Zhirui Hu, Xiaowo Wang

## Abstract

Functional genetic elements are one of the most essential units for synthetic biology. However, both knowledge-driven and data-driven methodology can hardly accomplish the complicated task of genetic elements design efficiently due to the lack of explicit regulatory logics and training samples. Here, we proposed a knowledge-constraint deep learning model named PccGEO to automatically design functional genetic elements with high success rate and efficiency. PccGEO utilized a novel “fill-in-the-flank” strategy with a conditional generative adversarial network structure to optimize the flanking regions of known functional sequences derived from the biological prior knowledge, which can efficiently capture the implicit patterns with a reduced searching space. We applied PccGEO in the design of *Escherichia coli* promoters, and found that the implicit patterns in flanking regions matter to the properties of promoters such as the expression level. The PccGEO-designed constitutive and inducible promoters showed more than 91.6% chance of success by in vivo validation. We further utilized PccGEO by setting a limited frequency of nucleotide modifications and surprisingly found that the expression level of *E. coli* sigma 70 promoters could show up to a 159.3-fold increase with only 10-bp nucleotide modifications. The results supported that the implicit patterns are important in the design of functional gene elements and validated the strong capacity of our method in the efficient design of functional genetic elements.

**Availability:** https://github.com/WangLabTHU/PccGEO

## 1. Introduction

Functional genetic regulatory elements are indispensable parts in synthetic biology.^1–5^ Traditional approaches for engineering genetic elements mainly rely on strong biological prior knowledge, such as transcription factor binding sites (TFBSs) and nucleosome disfavoring sequences, as these sequences are supposed to compose the kernel of *cis*-regulatory logics and thus be the key of gene regulation^6,7^. However, recent studies showed that the regulatory patterns of genetic elements are largely implicit depending on both the embedded sequences and their flanking regions. For example, de Boer et al. showed that 94% of gene expression were contributed by prevalent weak regulatory patterns which alter expression < 2-fold in eukaryotic promoters^8^. These prevalent weak patterns may refer to potential dependency between one motif and its flanking regions^9^, long range interactions between regions,^8,10^ or physicochemical properties limitations,^11–13^ Most of these weak patterns are too implicit to be summarized into concise design criteria. However, ignoring these implicit patterns would lead to a low rate of success in genetic elements design.^14^,^15^

Recently, data-driven learning methods offer potential solutions to capture complex relationships in multiple fields such as language modelling^16^ and images representation learning.^17,18^ Preliminary trials were also carried out to design genetic regulatory elements such as promoters,^19–21^ RNA switches, ^22^ guide RNA^23^,24 and ribosome-binding site (RBS),^25^ which suggested the great potential to employ data-driven methods, especially deep learning methods in biological sequence design. However, considering the exponentially growing biological sequence space and the limited training samples of functional elements, it is notoriously difficult to search for functional nucleotide combinations.^26^ Thus, proper methodologies combining both prior biological knowledge and data-derived implicit patterns should be carried out to narrow down the searching space for efficient genetic elements design.

Here, we proposed a “fill-in-the-flank” learning strategy to solve the problem. This strategy preserves the sequences derived from prior biological knowledge and then optimizes their flanking regions to endow implicit patterns of the whole regulatory element via conditional generative adversarial networks (cGANs).^27,28^ Attention-based methods^29^ were employed in the generator and discriminator of cGANs model to learn the long-range connection pattern. After training, sequences generated by the generator were further optimized by a genetic algorithm and a prediction model with a DenseNet structure.^30^

With the *Escherichia coli* promoter design task as an example, we demonstrated PccGEO’s capability to design the functional genetic elements both *in silico and in vivo*. By preserving the regions of –10 and –35 sequences, up to 100% of model-designed promoters showed higher expression levels compared with randomly generated promoters. Similarly, by preserving the *lacO* motif, up to 91.6% promoters with flanking region optimization retained the LacI-DNA interaction pattern and showed higher induced expression levels compared with the non-optimized group. We further utilized PccGEO to optimize sigma 70 promoters with limited nucleotide modification frequencies, and surprisingly found that genetic elements could show up to a 6.8-fold increase with only 5-bp nucleotide modifications, and 159.3-fold with 10-bp nucleotide modifications. In conclusion, the results support the implicit patterns are also the important factors in the genetic elements design and show the strong ability of our learning model for constructing well-characterized genetic elements.

## 2. Methods

### Problem definition

The target of our PccGEO model is to maximize the properties of functional elements, i.e. *max*_ŷ_ *prop*(ŷ), where ŷ denotes the genetic elements, and *prop*(⋅) denotes the property strength of elements, e.g., the gene expression level. Strong knowledge explicitly defines the annotated regions, and we use *x*_*prior*_to represent the annotated regions in the original element. In the *x*_*prior*^′^_, the annotated regions would be reserved, and other regions would be discarded. we defined *x*_*prior*_= {*m* _*i*_ ×*y*_*i*_}, *m* _*i*_ ∈{0, 1}. *m* is a binary vector, *y* is the natural element, and *i* means the position index. We use 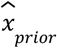 to denote the corresponding regions in ŷ. Then using strong knowledge constraints to design genetic elements ŷ, which should be depicted as:

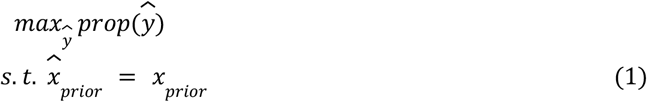

The above equation means the strong prior regions in ŷ and y should be the same. In the next two sections, we will demonstrate PccGEO model to design the functional genetic elements ŷ based on cGANs, DenseLSTM Predictor and genetic algorithm.

### Implicit patterns learning through “fill-in-the-flank” cGANs model

The implicit patterns would greatly influence the properties of our designed biological parts. These patterns shaped the flanking regions of the annotated regions corresponding to strong knowledge, e.g, complex interaction relationship between the annotated regions and flanking regions, and the interactions inside the flanking regions. To better utilize the prior knowledge and learn the implicit patterns, we defined a “fill-in-the-flank” strategy based on cGANs **(Fig. 1A**). Previous studies have demonstrated the preponderance of Generative adversarial networks (GANs) in designing multiple synthetic biological elements by learning their distributions in the latent space,^31^ which suggested the regulatory patterns could be learned by the GANs model. We follow the main structure of the generator and discriminator in Wang et al. work,^31^ but insert one attention based layer on the top of both generator and discriminator to better capture implicit patterns within flanking regions. Besides, “fill-in-the-flank” strategy instructs cGANs model to take the annotated regions as the model input, and learn to fill in the flanking regions by learning their implicit-pattern-decided distributions **(Fig. 1B**). The intuition behind the strategy is that the cooperation between the annotated regions and flanking regions is important for the regulation, such as the interaction between TFBSs and flanking DNA shape^11^ would affect TF binding. Thus, more accurate implicit patterns could be learned by adding the “fill-in-the-flank” strategy.

**Fig. 1.**
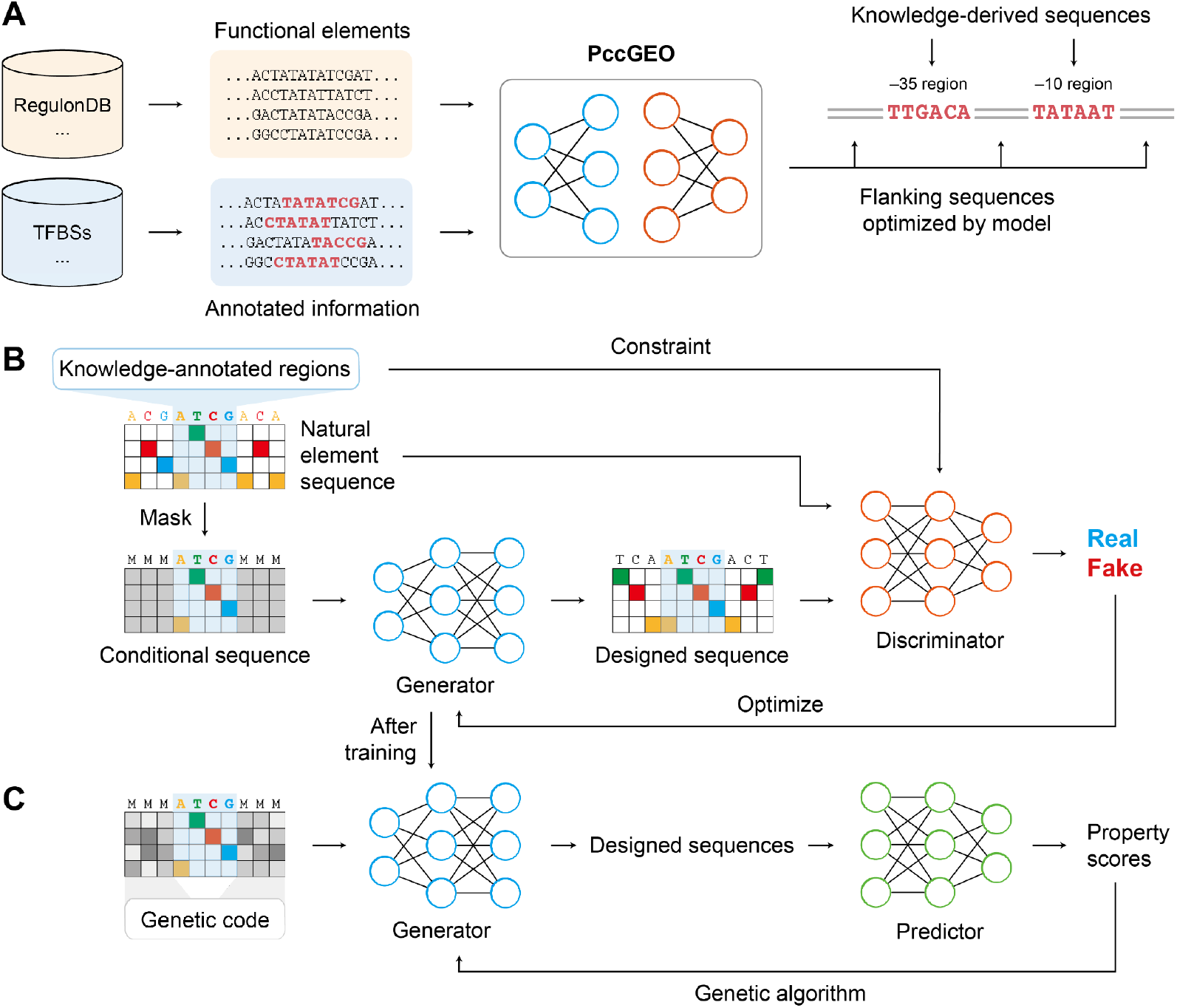
The landscape of our AI model. **A**, by combining the prior biological knowledge and the element sequences data, our model could automatically design functional elements. RegulonDB^32^ is a database containing the *E. coli* element sequences. TFBSs: transcription factor binding sites. **B**, the learning procedure of the cGAN, by taking the conditional sequence as the model input, cGANs could automatically design the functional sequences. **C**, the learning procedure of element optimization. with a prediction model, we optimize the design sequences by genetic algorithm.

The loss function of cGANs can be described as below:

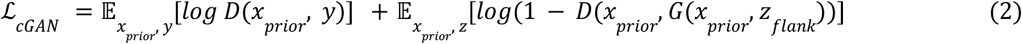

Where *Z*_*flank*_ denotes the latent codes of flanking regions, *G* and *D* represent the generator and discriminator respectively. To preserve the annotated regions such as TFBSs as strong constraints and force the cGANs model to learn the implicit patterns between the annotated and flanking regions, we added one L1 loss inspired by previous works in image generation^28^. This loss can be written as:

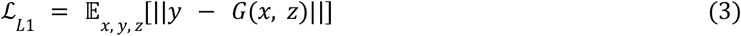

The final optimizing object in implicit patterns learning is:

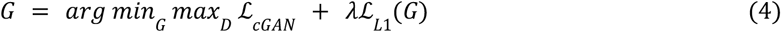

With the setting of ‘fill-in-the-flank’ task and the power of cGANs, generator captured implicit patterns information provides the basis for more efficient optimization.

### Accurate prediction offers guidance for properties optimization

Accurate predictions of target properties are crucial in our approach to optimize the flanking regions based on learned implicit patterns. Therefore the predictors should also take the implicit patterns into consideration, such as the long range interactions. Predictions of biological properties often suffer from the overfitting problem, thus the predictor should be carefully designed. We used a deep learning model for prediction. In the first layer, we used 1d convolutional kernels with 64 output channels to capture motif information to avoid the overfitting. Then we added the Long Short Term Memory (LSTM)^33^ architecture to capture the relationships between motifs. In order to extract potential long range relationship factors leading to the target properties, we adopted the DenseNet^30^ architecture to efficiently improve the network depth. We set 4 Dense Blocks with 2, 2, 4 and 2 Dense Layers in each block. We set the growth rate to 32. Each Dense Layer contains two convolutional layers with the kernel size as 1 and 3 respectively. Finally we added one fully connected layer to predict the property of interests. This predictor is denoted as *P*_*DenseLSTM*_.

Then we leveraged the genetic algorithm (GA) to optimize the generated flanking regions by maximizing the output of the predictor **(Fig. 1C**). We used the GA^34^ module in the sko python package to implement the genetic algorithm. The optimization process using the genetic algorithm from step t to t+1 could be written as:

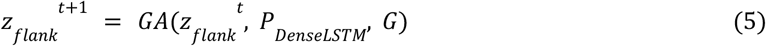

The restrictions of “fill-in-the-flank’’ is that it requires the strong prior-knowledge-decided annotated regions given to the model input.

### Polishing elements through identification of optimization positions

To enable a wider application, we also consider the task of optimizing existing elements without any prior knowledge. We extended our “fill-in-the-flank” strategy to the case where the model detected the optimized regions automatically. The constraint on the number of modified positions *N*_*polish*_ is added for utilizing the existing useful relationships and reducing the searching space. Therefore this could be viewed as one polish task to optimize specific biological parts with limited modification. The genetic algorithm simultaneously optimizes two parts: the discrete flanking region position vector *pos*_*flank*_ and the continuous latent codes of flanking regions *z* _*flank*_. Each component of the flanking position vector represents one modified position index, therefore we satisfied the modification limitation by controlling the length of the modification positions vector. This can be formulated as:

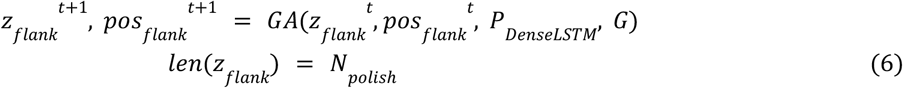

We set the size of populations in the genetic algorithm to 12*1024 and the max iteration number to 100. We used the “vectorization” mode in the sko package to facilitate the optimization. The mutation probability was set to 0.005.

## 3. Experiments setup

### *In silico* evaluation experiments

*In silico* evaluation was performed to test whether our method successfully improved the target property’s strength. Three experiments were carried out, including optimizing the expression level of constitutive promoters with –10 and –35 regions preserved, inducible promoters with –10, –35 regions preserved and one additional operator inserted, and the natural promoters with 10 bps modification constraints. We seperated the datasets into 3 folds, we used 2 folds to train the predictor for the genetic algorithm and used the other fold to train another validation predictor for evaluation. The original promoters were selected from the fold corresponding to the validation predictor.

### *In vivo* evaluation experiments

We also carried out 3 experiments in *E. coli*. We first fixed –10 and –35 regions with 5 kinds of combinations of identified sequences. Then we used our method to improve the promoter activity. Secondly, we preserved –10 and –35 regions of natural promoters and inserted one operator, and other regions were treated as flanking regions to be optimized. We compared our designs with the promoters directly inserted by one operator. Finally we optimized promoters within the RegulonDB with 5 bps modification and 10 bps modification.

### Experimental dataset for model training

Two types of RNA-seq experimental datasets were used in the training process. The 165 bp RNA-seq dataset contained a total of 29249 regulatory sequences from 184 prokaryotic genomes. The regulatory sequence was defined as 165 bp upstream of the gene start codon. The regulatory library was cloned into a p15A vector and transformed into *E. coli* MG1655.^35^ Another RegulonDB dataset contained a total of 14,868 transcriptional start sites (TSS) candidates from the *E. coli* MG1655 genome. The promoter sequence was defined as 100 bp upstream of the TSS.^36^

### Bacterial Strains and Plasmids

All transformations and fluorescence assays were performed in the *E. coli* strain trans5α [F-, φ80d, *lac*ZΔM15, Δ(*lac*ZYA-*argF*), U169, *end*A1, *rec*A1, *hsd*R17(r_k_^-^, m_k_^+^), *sup*E44λ-, *thi*-1, *gyr*A96, *rel*A1, *pho*A]. All plasmid constructions were carried out using NEBuilder HiFi DNA assembly reaction according to standard Gibson assembly. The activity of the promoter was measured by the intensity of expressed sfGFP. Promoter constructions were cloned into the vector pTPR with a p15A origin of replication and chloramphenicol resistance. A terminator DT5 was inserted at the 5’ end of the promoter to avoid the influence of the upstream sequence, and the insulator Riboj was inserted at the 3’end of the promoter to ensure that the same transcript was produced. The ribosomal binding site sequence BBA_B0034 is used at the 5’ end of *sfgfp gene*. Double terminators BBa_B0015 were used to terminate *sfgfp* gene transcription. For inducible promoter constructs, an additional LacI protein expression cassette was inserted at the 5’ end of terminator DT5 in the opposite reading direction.

### Assay of promoter strength

The strain containing the target promoter plasmid was cultured overnight (16 hours) in 5ml LB medium supplemented with 50 ug/ml Chloramphenicol at 37°C in a shaker at 220 rpm for promoter activity validation. The overnight cultures were diluted 1:100 into fresh LB medium supplemented with 50 ug/ml Chloramphenicol in triplicate. For inducible promoters, a final concentration of 0.1 mM IPTG was added into the medium when diluted. After incubating for another 6 hours, 150ul of cultures was added to a flat-bottomed 96-well microplate (Corning 3603) and repeated the measurements of the optical density at 600 nm (OD_600_) and fluorescence (relative fluorescence units [RFU]; excitation at 485 nm and emission at 520 nm) with the Varioskan Flash (Thermo). The background fluorescence was measured using the 150 ul fresh LB medium and a strain harboring a promoterless plasmid. The strength of the promoter is defined as the average of fluorescence/OD_600_ after subtracting background fluorescence.

## 4. Results

### *In silico* evaluation of PccGEO verified the biological parts transferring ability

We first evaluated PccGEO *in silico* on optimizing the expression levels by preserving –10 and –35 regions. Then we added another operator inserted. We used 3-fold cross validation *in silico*. We observed our method could successfully optimize most promoters in the perspective of validation predictors. The high success rate could be attributed to the combination of the strong knowledge and the implicit pattern learning **(Fig. 2A**). We then evaluated on optimizing existing promoters with 10 bps modification. We observed that all the validated promoters’ expression levels were improved. We then embedded all promoters into the low dimension space using the unsupervised representation approach DeepInfoMax ^37^ for more comprehensive investigation. We used UMAP^38^ to plot promoter embeddings in the 2 dimension space by utilizing the output of the penultimate layer in the DenseLSTM predictor. We could clearly see promoters with low and high expression levels located on left and right sides, which could imply promoters with different expression levels had different distributions. We observed the tendency of our design moving towards the regions corresponding to high expression levels (**Fig. 2B**). For promoters with limited modification numbers, even though the Hamming distances between original and designed promoters are very close, we could observe their function embeddings are moving from low expression regions to high expression regions. (**Fig. 2B**). These experiments computationally demonstrated our method’s success with the assistance of strong prior knowledge and learning the implicit patterns.

**Fig. 2.**
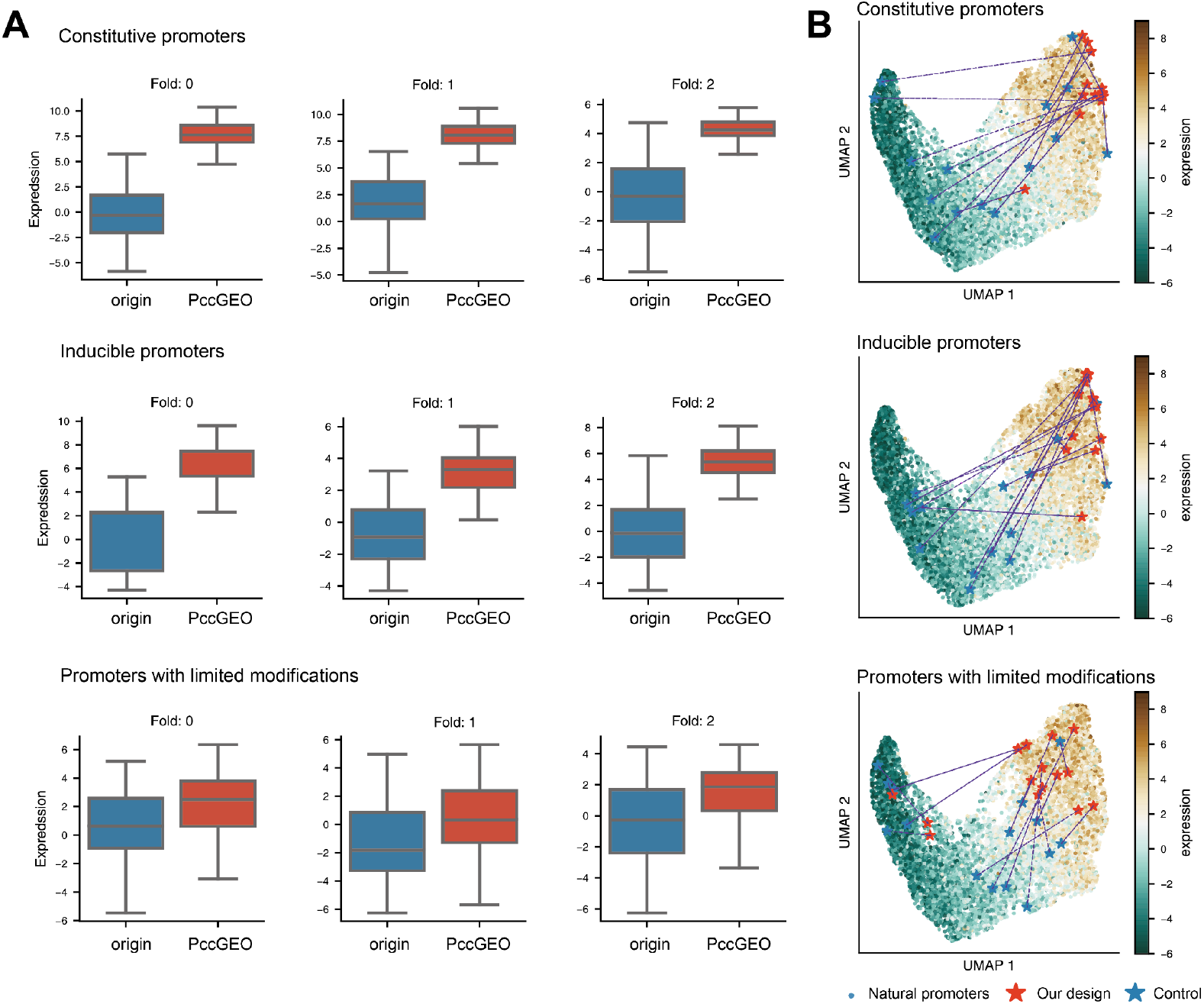
*In silico* promoter optimization results. **A**, the *in silico* evaluations, from top to bottom, the figures are results of preserving –10 and –35 regions, preserving –10, –35 and operator regions of inducible promoters, and optimization with limited modification numbers. **B**, the promoters in **A** plot in the embeddings spaces, each dot represents one natural promoter; blue green means the high expression level; red stars and blue stars mean the original promoters and promoters after our design, dashed lines connect the corresponding pairs.

### Implicit patterns matter to the gene expression could be learned by PccGEO

The previous studies have mentioned the regulatory pattern that is rather implicit depending on both the embedded motif sequences and their flanking regions. We illustrated one of the implicit patterns, DNA shape, matters to the gene expression. DNA shape ^11^ is a kind of physicochemical properties of DNA sequence which define the structural information, including minor groove width, propeller twist, etc **(Fig. 3A**). We predicted the DNA shape feature of natural promoter sequences from the Monte Carlo approach^39^ and used UMAP^38^ to plot DNA shape embeddings into the two dimension space. We could find that the high and low expression sequences could be separated into the left and right side of the space, which illustrated that DNA shape as the implicit pattern would matter to the gene expression. We optimized 15 promoters by the PccGEO model, and observed half of them were directly moved from the right to the left **(Fig. 3B**), which demonstrated that PccGEO could learn the DNA shape pattern in the genetic elements design.

**Fig. 3.**
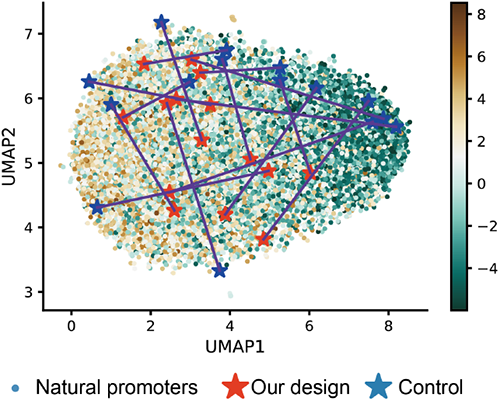
Dimension reduction of DNA shapes feature of promoter sequence. We showed the optimized trajectory of 15 natural promoter sequences. Red stars and blue stars mean the original promoters and promoters after our design, dashed lines connect the corresponding pairs.

### The flanking sequences optimization enhances the constitutive promoter in *E. coli*

After successfully improving the expression of the promoter *in silico*, biological experiments were carried out to evaluate the performance of constitutive promoters. In *E. coli*, transcription requires the synergy of RNA polymerase, σ factors, and promoter sequences. σ factors interact with an RNA polymerase core enzyme to form the RNAP holoenzyme and direct the complex to promoter regions by recognizing specific regulatory sequences^40^. Previous studies have found that the structures of highly expressed constitutive promoters have common features in *E. coli*, such as –10 and –35 elements and their spacing constraints^41^. We chose five combinations of –10 and –35 elements and the length of the spacer between –10 and –35 elements was specified to be 17 bp which was proper for *E. coli* promoters (**Fig. 4A**). Strong constraints in constitutive promoter design were the sequences of –10 and –35 elements and the length of the spacer between –10 and –35 elements, and our model optimized the flanking sequence to improve the strength of promoters. The top two predicted promoters from the genetic algorithm were chosen to promote the expression of sfGFP in *E. coli*. Two 165 bp promoters with the same –10 and –35 regions and randomly generated flanking regions were employed as negative controls. The results indicated that 100% model-designed promoters showed high expression levels compared with random generated sequence (t-test with Benjamini Hochberg correction, FDR < 5%) (**Fig. 4B-F**). Due to the canonical –10 (TATAAT) and –35 (TTGACA) combination having a high affinity with sigma factors, the control groups also showed a high expression level, but the variation within the group was large, which revealed the implicit patterns played a key role even the prior knowledge is very strong. The finite improvement of the canonical –10 and –35 combination group could be possibly owed to the limited cell production capacity (**Fig. 4B**). For non-canonical –10 and –35 combination features, model-designed promoters showed great improvement of expression level compared with the control group (**Fig. 4C-F**). The experiment results of constitutive promoters proved that our model could improve the promoter expression strength through flanking sequence optimization and the introduction of prior knowledge of functional regions.

**Fig. 4.**
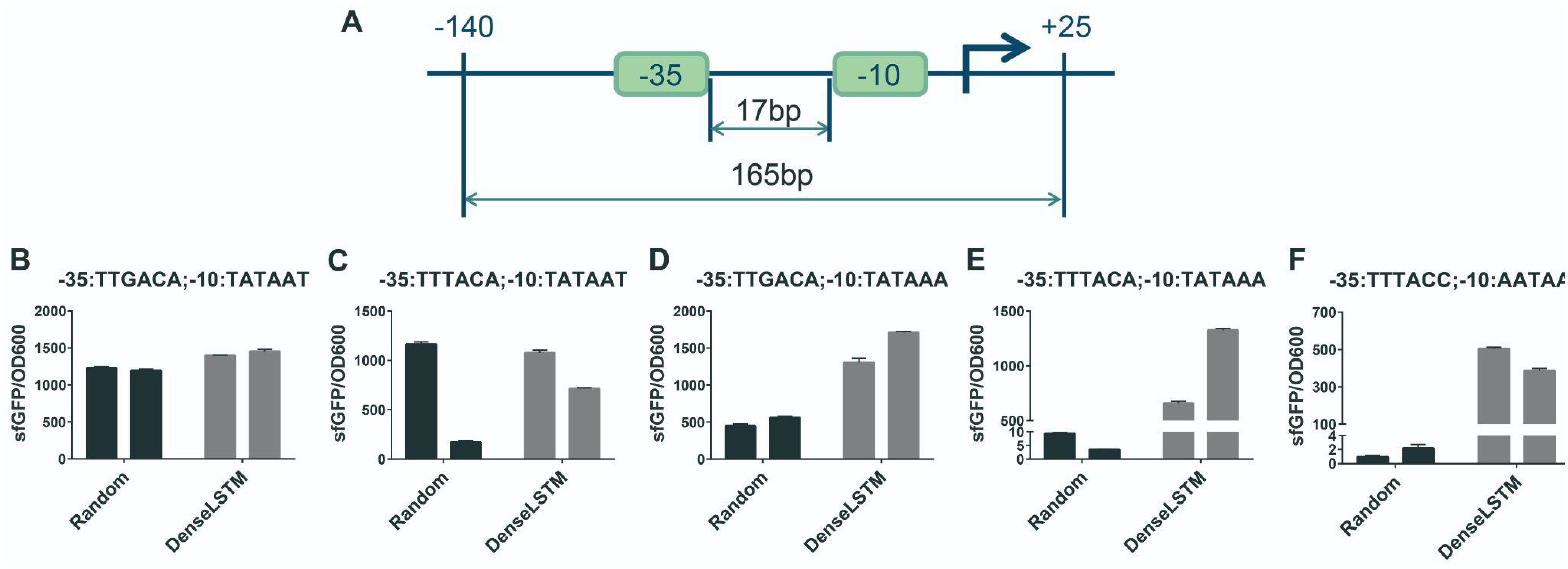
Design and experimental results of constitutive promoters in *E. coli*. **A**, the structure of constitutive promoter used in *E. coli*. We preserved –10 and –35 regions as the annotated regions in the sequences. **B-F**, the expression levels of constitutive promoters composed by five combinations of –10 and –35 motifs respectively. Random group: the flanking sequences generated randomly; DenseLSTM group: the flanking sequences generated by our model and filtered by the DenseLSTM predictor. Values and error bars represent the mean and the s.d. (n=3).

### Strong knowledge driven inducible promoters design with implicit patterns

Except housekeeping genes are always active, most genes are only expressed in specific environments. The inducible promoter only activates the transcription under specific circumstances, like the most classic example with *lac* operon in *E. coli*. The upstream sequence of *lac* operon expresses the LacI protein, which blocks RNAP from binding to the operator. This inhibition can be relieved by the addition of lactose and its analogues^42^. Employing the LacI-DNA interaction pattern as strong prior, a series of inducible promoters were designed based on the promoter sequence from the dataset (**Fig. 5A**). The *lacO* element (17 bp) was inserted at 15 bp downstream from the –10 motif as the LacI protein binding sequence, which has been proven to be the proper position of *lac* promoter regulation by previous studies^43^. We randomly selected 5 promoter sequences from the 165 bp RNA-seq dataset as the initial sequence. After analyzing each promoter sequence, the –10 motif, –35 motif, the length of the spacer, the position and sequence of *lacO* were kept as strong prior constraints. The flanking regions were designed by our model to improve the strength of the promoter in inducible promoter design. The control group for each promoter is set by substituting its 17 bp sequence downstream of the +5 region of the original promoter with the *lacO* sequence (**Fig. 5B**). Due to the lack of learning samples for fold-change information, we only focused on the optimization of their expression levels. The top 3 promoters predicted by DenseLSTM were chosen to measure the expression level of sfGFP. The expression of the control group promoter was reduced compared with the original one under induction because of the destruction of implicit patterns, e.g., potential interactions or improper flanking sequence. On the contrary, 91.6% of the inducible promoters designed by our model showed a higher expression level than the original promoter under induction (t-test with Benjamini Hochberg correction, FDR < 5%). And the introduction of IPTG increased the expression level of the designed-promoters illustrating that the function of *lacO* motif has been successfully retained (**Fig. 5C-F**).

**Fig. 5.**
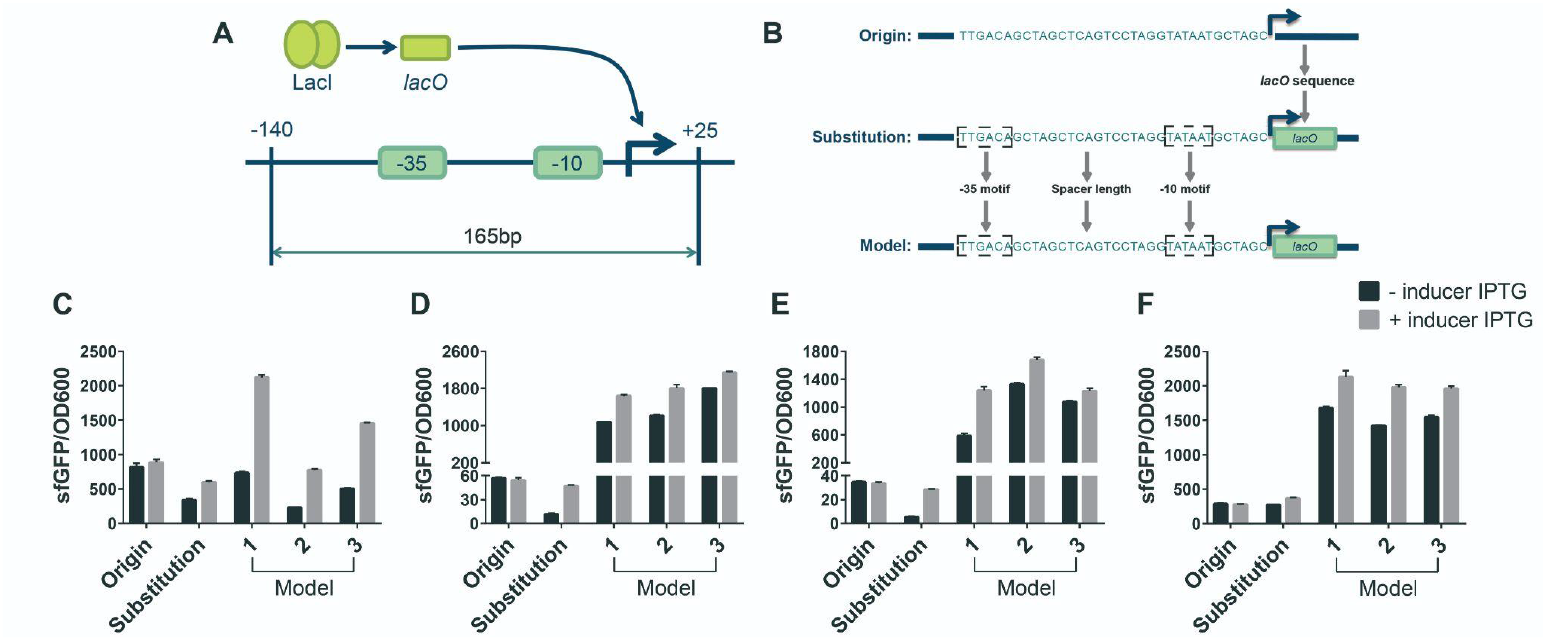
Design and experimental results of inducible promoters in *E. coli*. **A**, the structure of designed inducible promoters with *lacO* sequence based on 165bp RNA-seq dataset promoter in *E. coli*. **B**, the structure diagram of origin, substitution and model promoter. Origin group: the promoter sequence was randomly selected in 165bp RNA-seq dataset; Substitution group: the *lacO* sequence was inserted at 15 bp downstream of -10 motif; Model group: the flanking sequence optimization by our model after preserving the prior sequence. **C-F**, the expression level of promoters with or without inducer IPTG. Values and error bars represent the mean and the s.d. (n=3).

### Promoter expression optimization within limited nucleotides modification

The functional regulatory elements design based on strong prior knowledge have been verified in constitutive and inducible promoters experiments. It is a common but difficult situation that regulatory elements with insufficient information are needed to be optimized, such as cross-species elements transfer and new regulatory elements application. For those regulatory elements, our model could automatically select the discordant sequence regions as the target optimization regions which have been proven *in silico*.

Here we chose six promoter sequences from the RegulonDB database for optimization and verified the performance of our model through biological experiments. Nucleotides modification with 5 or 10 base pairs was added to show the performance of the model. The limitation of nucleotides number would prevent the excessive destruction of initial sequence relationships. The DenseLSTM predictor was used to evaluate the expression of optimized promoter sequences, and the top 3 promoters were chosen to drive the expression of sfGFP in *E. coli*. Compared with the initial promoter sequence, model-designed promoters indicated up to a 6.8-fold increase with only 5-bp nucleotide modifications, and 159.3-fold increase with 10-bp modification on average. Moreover, the expression level of the 10 bp modification group in 83.3% of optimization examples was significantly higher than the 5 bp modification group (t-test with Benjamini Hochberg correction, FDR < 5%). Therefore our model could improve the expression level of a promoter by modifying limited nucleotides (**Fig. 6**).

**Fig. 6.**
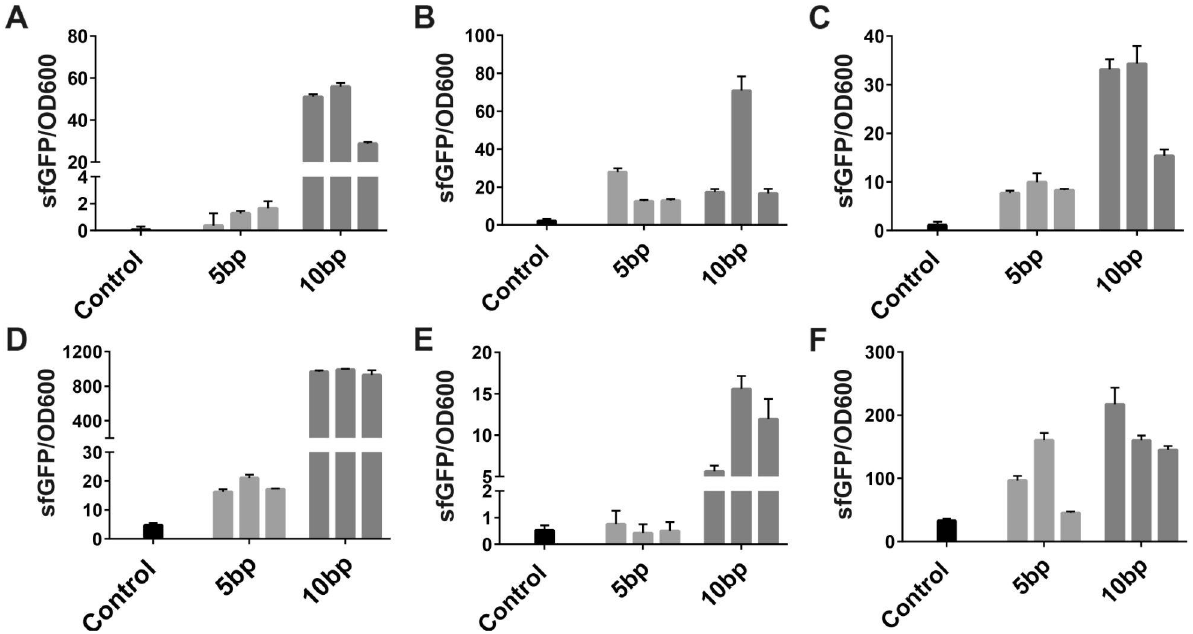
The expression levels of promoters with limited nucleotides modification. **A-F**, the expression level of six optimization experiments of different initial promoter sequences. Control group: promoter sequence randomly selected in the RegulonDB database; 5bp group and 10bp group: 5 or 10 base pairs nucleotide modifications on the initial promoter sequence, and the top 3 promoters in the DenseLSTM predictor were used. Values and error bars represent the mean and the s.d. (n=3).

## 5. Discussion

In this study, a prior knowledge based and implicit patterns attentioned learning framework named PccGEO was proposed to automatically design functional genetic elements. PccGEO utilized cGANs model combined with “fill-in-the-flank” strategy to efficiently capture the implicit patterns in the flanking regions, and an accurate DenseLSTM prediction model to offer guidance for properties optimization. PccGEO could achieve 100% and 91.4% chance of success in the constitutive and inducible promoter design in *E*.*coli*, which verified the high efficiency of flanking regions optimization. Interestingly, we further found that our model could automatically optimize the properties of functional elements with limited nucleotides modification, which could achieve 6.8-fold gene expression increase with 5-bp modification, and 159.3-fold increase with 10-bp modification. These results supported that implicit patterns were important for gene regulation, and our model could utilize them to design genetic elements in an efficient way.

Admittedly, our work still could be further improved. Although we know that the implicit pattern has a significant influence on functional elements, the black box characteristics of deep neural networks hinder the understanding of such implicit patterns. But we still have shown that one of the implicit patterns - DNA shape could influence the gene expression, and have shown our PccGEO model could learn DNA shape to optimize the genetic elements. Besides, for the inducible promoter design task, the optimization of induction rates is more important. However, due to the lack of a database of inducible promoter sequences in the flanking regions, the only thing we could do is to optimize the expression strength based on our PccGEO model. Future work will be done to optimize induction rates by generating experimental data with massive parallel reporter assay^44^. Furthermore, more experiments will be done to prove that our method could be generalized to the eukaryotic genetic elements design, to capture the more complex regulatory rules in both annotated and unannotated regions in the functional elements.

In conclusion, we provided a novel method to design functional genetic elements based on both the prior biological knowledge and the implicit patterns learning methods. By utilizing a “fill-in-the-flank” learning strategy and carefully designed learning model, the functional constitutive and inducible synthetic promoters were designed by our model, and their properties were validated *in silico* and *in vivo*. Our work provided a new insight into a new genetic elements design methodology by incorporating biological knowledge as well as other implicit patterns through data learning to efficiently explore the sequence space and raise the design success rate.

## Reference

1 de Lange O, Klavins E, Nemhauser J. Synthetic genetic circuits in crop plants. Curr Opin Biotechnol 2018;49:16–22.

2 McNerney MP, Doiron KE, Ng TL, Chang TZ, Silver PA. Theranostic cells: emerging clinical applications of synthetic biology. Nat Rev Genet 2021;22:730–46.

3 Lynch SA, Gill RT. Synthetic biology: new strategies for directing design. Metab Eng 2012;14:205–11.

4 Ye W, Haochen W, Minghao YAN, Guanhua HU, Xiaowo W. Design of biomolecular sequences by arti?cial intelligence. Synthetic Biology Journal 2021;2:1.

5 Tianying Chen, Xue Zhang, Qiong Wu. Recent progress in research and application of engineered implanted cells for biomedical applications. Quant Biol 2021;0:0.

6 Weingarten-Gabbay S, Nir R, Lubliner S, Sharon E, Kalma Y, Weinberger A, et al. Systematic interrogation of human promoters. Genome Res 2019;29:171–83.

7 Mogno I, Kwasnieski JC, Cohen BA. Massively parallel synthetic promoter assays reveal the in vivo e?ects of binding site variants. Genome Res 2013;23:1908–15.

8 de Boer CG, Vaishnav ED, Sadeh R, Abeyta EL, Friedman N, Regev A. Deciphering eukaryotic gene-regulatory logic with 100 million random promoters. Nat Biotechnol 2020;38:56–65.

9 Liu X, Gupta STP, Bhimsaria D, Reed JL, Rodríguez-Martínez JA, Ansari AZ, et al. De novo design of programmable inducible promoters. Nucleic Acids Res 2019;47:10452–63.

10 Yu TC, Liu WL, Brinck MS, Davis JE, Shek J, Bower G, et al. Multiplexed characterization of rationally designed promoter architectures deconstructs combinatorial logic for IPTG-inducible systems. Nat Commun 2021;12:325.

11 Rohs R, West SM, Sosinsky A, Liu P, Mann RS, Honig B. The role of DNA shape in protein–DNA recognition. Nature 2009;461:1248–53.

12 Mathelier A, Xin B, Chiu T-P, Yang L, Rohs R, Wasserman WW. DNA Shape Features Improve Transcription Factor Binding Site Predictions In Vivo. Cell Syst 2016;3:278–86.e4.

13 Zhou T, Shen N, Yang L, Abe N, Horton J, Mann RS, et al. Quantitative modeling of transcription factor binding speci?cities using DNA shape. Proc Natl Acad Sci U S A 2015;112:4654–9.

14 Vogl T, Ruth C, Pitzer J, Kickenweiz T, Glieder A. Synthetic core promoters for Pichia pastoris. ACS Synth Biol 2014;3:188–91.

15 Carr SB, Beal J, Densmore DM. Reducing DNA context dependence in bacterial promoters. PLoS One 2017;12:e0176013.

16 Devlin J, Chang M-W, Lee K, Toutanova K. BERT: Pre-training of Deep Bidirectional Transformers for Language Understanding. arXiv [csCL] 2018.

17 He K, Zhang X, Ren S, Sun J. Deep residual learning for image recognition. And Pattern Recognition 2016.

18 Simonyan K, Zisserman A. Very Deep Convolutional Networks for Large-Scale Image Recognition. arXiv [csCV] 2014.

19 Van Brempt M, Clauwaert J, Mey F, Stock M, Maertens J, Waegeman W, et al. Predictive design of sigma factor-speci?c promoters. Nat Commun 2020;11:5822.

20 Hossain A, Lopez E, Halper SM, Cetnar DP, Reis AC, Strickland D, et al. Automated design of thousands of nonrepetitive parts for engineering stable genetic systems. Nat Biotechnol 2020;38:1466–75.

21 Kotopka BJ, Smolke CD. Model-driven generation of arti?cial yeast promoters. Nat Commun 2020;11:2113.

22 Angenent-Mari NM, Garruss AS, Soenksen LR, Church G, Collins JJ. A deep learning approach to programmable RNA switches. Nat Commun 2020;11:5057.

23 Chuai G, Ma H, Yan J, Chen M, Hong N, Xue D, et al. DeepCRISPR: optimized CRISPR guide RNA design by deep learning. Genome Biol 2018;19:80.

24 Wang D, Zhang C, Wang B, Li B, Wang Q, Liu D, et al. Optimized CRISPR guide RNA design for two high-?delity Cas9 variants by deep learning. Nat Commun 2019;10:4284.

25 Ding N, Yuan Z, Zhang X, Chen J, Zhou S, Deng Y. Programmable cross-ribosome-binding sites to ?ne-tune the dynamic range of transcription factor-based biosensor. Nucleic Acids Res 2020;48:10602–13.

26 Linder J, Bogard N, Rosenberg AB, Seelig G. A Generative Neural Network for Maximizing Fitness and Diversity of Synthetic DNA and Protein Sequences. Cell Syst 2020;11:49–62.e16.

27 Mirza M, Osindero S. Conditional Generative Adversarial Nets. arXiv [csLG] 2014.

28 Isola P, Zhu J-Y, Zhou T, Efros AA. Image-to-Image Translation with Conditional Adversarial Networks.

29 Vaswani A, Shazeer N, Parmar N, Uszkoreit J, Jones L, Gomez AN, et al. Attention Is All You Need.

30 Huang G, Liu Z, Van Der Maaten L, Weinberger KQ. Densely Connected Convolutional Networks.

31 Wang Y, Wang H, Wei L, Li S, Liu L, Wang X. Synthetic promoter design in Escherichia coli based on a deep generative network. Nucleic Acids Research 2020:6403–12. https://doi.org/10.1093/nar/gkaa325.

32 Santos-Zavaleta A, Salgado H, Gama-Castro S, Sánchez-Pérez M, Gómez-Romero L, Ledezma-Tejeida D, et al. RegulonDB v 10.5: tackling challenges to unify classic and high throughput knowledge of gene regulation in E. coli K-12. Nucleic Acids Res 2019;47:D212–20.

33 Hochreiter S, Schmidhuber J. Long short-term memory. Neural Comput 1997;9:1735–80.

34 Whitley D. A genetic algorithm tutorial. Stat Comput 1994;4:65–85.

35 Johns NI, Gomes ALC, Yim SS, Yang A, Blazejewski T, Smillie CS, et al. Metagenomic mining of regulatory elements enables programmable species-selective gene expression. Nat Methods 2018;15:323–9.

36 Thomason MK, Bischler T, Eisenbart SK, Förstner KU, Zhang A, Herbig A, et al. Global transcriptional start site mapping using di?erential RNA sequencing reveals novel antisense RNAs in Escherichia coli. J Bacteriol 2015;197:18–28.

37 Devon Hjelm R, Fedorov A, Lavoie-Marchildon S, Grewal K, Bachman P, Trischler A, et al. Learning deep representations by mutual information estimation and maximization. arXiv [statML] 2018.

38 McInnes L, Healy J, Saul N, Großberger L. UMAP: Uniform Manifold Approximation and Projection. Journal of Open Source Software 2018:861. https://doi.org/10.21105/joss.00861.

39 Zhou T, Yang L, Lu Y, Dror I, Dantas Machado AC, Ghane T, et al. DNAshape: a method for the high-throughput prediction of DNA structural features on a genomic scale. Nucleic Acids Res 2013;41:W56–62.

40 Feklístov A, Sharon BD, Darst SA, Gross CA. Bacterial sigma factors: a historical, structural, and genomic perspective. Annu Rev Microbiol 2014;68:357–76.

41 Campbell EA, Muzzin O, Chlenov M, Sun JL, Anders Olson C, Weinman O, et al. Structure of the Bacterial RNA Polymerase Promoter Specificity σ Subunit. Molecular Cell 2002:527–39. https://doi.org/10.1016/s1097-2765(02)00470-7.

42 Rezniko? WS. The lactose operon-controlling elements: a complex paradigm. Molecular Microbiology 2006:2419–22. https://doi.org/10.1111/j.1365-2958.1992.tb01416.x.

43 Lewis M. The lac repressor. C R Biol 2005;328:521–48.

44 Melnikov A, Murugan A, Zhang X, Tesileanu T, Wang L, Rogov P, et al. Systematic dissection and optimization of inducible enhancers in human cells using a massively parallel reporter assay. Nat Biotechnol 2012;30:271–7.

